# Response expectations shape serial dependence and stimulus processing

**DOI:** 10.1101/2025.10.22.683912

**Authors:** Junlian Luo, David Pascucci

**Affiliations:** Psychophysics and Neural Dynamics Lab, Department of Radiology, Lausanne University Hospital, (CHUV) and University of Lausanne (UNIL), Lausanne, Switzerland; The Sense Innovation and Research Center, Lausanne, Switzerland

**Keywords:** serial dependence, inverted encoding, EEG, perceptual decision-making

## Abstract

Perceptual decisions are biased by recent history, yet the balance between attractive and repulsive effects varies across contexts. Here, we tested whether trial-by-trial response expectations shape the direction of history biases in serial dependence during orientation reproduction. Behaviorally, we found that no-response trials—especially when rare—reduced attractive biases and enhanced repulsive biases. EEG results revealed stronger evoked responses and amplified neural representations for stimuli following no-response trials. Together, these findings suggest that interrupting the perception–action cycle fosters a state of re-engagement with current input and disengagement from past stimuli, indicating that serial dependence is a flexible process dynamically modulated by task expectations and transient shifts in sensory processing.

## Introduction

Perceptual decisions are shaped not only by current sensory input but also by the temporal structure and history of recent events. A well-known example is *serial dependence*, in which decisions about the current stimulus are biased by stimuli encountered a few trials before ^1^.

Over the past decade, numerous studies have documented serial dependence across a wide range of visual tasks. These studies typically reported *attractive* biases, whereby visual decisions are biased toward prior stimuli ^see 2,3 for reviews^. A classic case is the orientation adjustment task: when reproducing the orientation of the current stimulus, particularly with weak, uncertain, and brief stimuli ^4–7^, observers make systematic errors in the direction of the orientations presented on recent trials. Similar effects have been observed for other basic features such as motion and color, as well as for more complex stimuli such as facial expressions, emotions, or even aesthetic preferences ^8–11^, suggesting that serial dependence reflects a general characteristics of human perception and cognition ^12^.

Despite its pervasiveness across modalities and even species ^13,14^, the magnitude and direction of serial dependence vary considerably across individuals and task conditions ^3,15–17^. In some cases, serial dependence is dominated by *repulsive* biases—perceptual reports are biased away from previous stimuli—and overall behavioral data suggest that attractive and repulsive components coexist and interact at the individual level ^15,18–24^. These two opposite forms of serial dependence are thought to arise from influences of prior events at different processing stages ^18,25,26^, yet the conditions under which each dominates remain unclear.

One factor proposed to influence this balance is the continuity of perception–action cycles. Most studies reporting attractive biases involve sequences of stimuli followed by overt responses. Manipulations of response requirements, however, show that attraction is reduced—or can even reverse into repulsion—following trials in which no response was required ^15,18,27,28^. For example, in orientation adjustment tasks, observers typically show attractive serial dependence after runs of response trials, whereas the same task can yield repulsive biases when the immediately preceding trial required no response ^15,18^. This pattern suggests that response requirements exert at least a modulatory effect on how previous stimuli influence current decisions, but the underlying mechanisms remain debated ^1,18,27–30^.

We hypothesized that the modulatory effects of prior response requirements reflect trial-by-trial expectations that shape processing of the current stimulus and, consequently, the influence of past events. In paradigms manipulating response requirements, participants may anticipate a response trial following a no-response trial—either due to perceived regularities in alternating events ^31^ or simply because response trials typically occur more frequently than no-response trials ^1,18,27^. This expectation to deliver a response could lead to a re-engagement with the task and stimuli, thereby reducing the influence of prior trials or even promoting a stronger tendency to discriminate the current stimulus from previous ones, potentially resulting in repulsive biases. In other words, interrupting the perception–action cycle may induce a state of stronger engagement with the current stimulus and relative disengagement from past events.

We tested this hypothesis using behavioral and EEG data. In a first behavioral experiment, we manipulated the ratio of response to no-response trials across blocks of an orientation adjustment task. The rationale is that if no-response trials induce an expectation of an upcoming response, this effect should depend on response frequency: the rarer the no-response trials, the stronger the expectation following them, and consequently, the greater the modulation of serial dependence. In a second experiment, we analyzed EEG data from a task in which response and no-response trials occurred with equal probability. We examined overall changes in EEG activity and in stimulus representations to identify neural correlates of the modulation of serial dependence following no-response trials.

Across both experiments, serial dependence patterns were systematically altered after no-response trials, showing reduced attraction and increased repulsion. This effect was evident when no-response trials were rare (Experiment 1) and also when they were equally likely as response trials (Experiment 2). In the latter case, EEG analyses revealed neural signatures indicative of increased attentional engagement and enhanced stimulus processing after no-response trials, consistent with the behavioral reduction in attraction and increase in repulsion. Together, these findings suggest that trial-by-trial expectations about response requirements modulate serial dependence: no-response trials appear to trigger a transient reset or re-engagement process that shifts the weighting of past versus current sensory information. These results indicate that serial dependence is not a fixed property of perceptual decision-making but a flexible phenomenon shaped by cognitive state and task context, offering new opportunities to relate behavioral variability and individual differences to fluctuations in internal states.

## Results

### Experiment 1

Participants performed an orientation adjustment task under three response-frequency conditions, run in separate blocks: a low-response condition (25% response trials), an equal-response condition (50%), and a high-response condition (75%). In no-response trials, participants were presented with a neutral stimulus (a circular frame without response cues, Figure 1A) and instructed to passively wait for the next trial.

**Figure 1.**
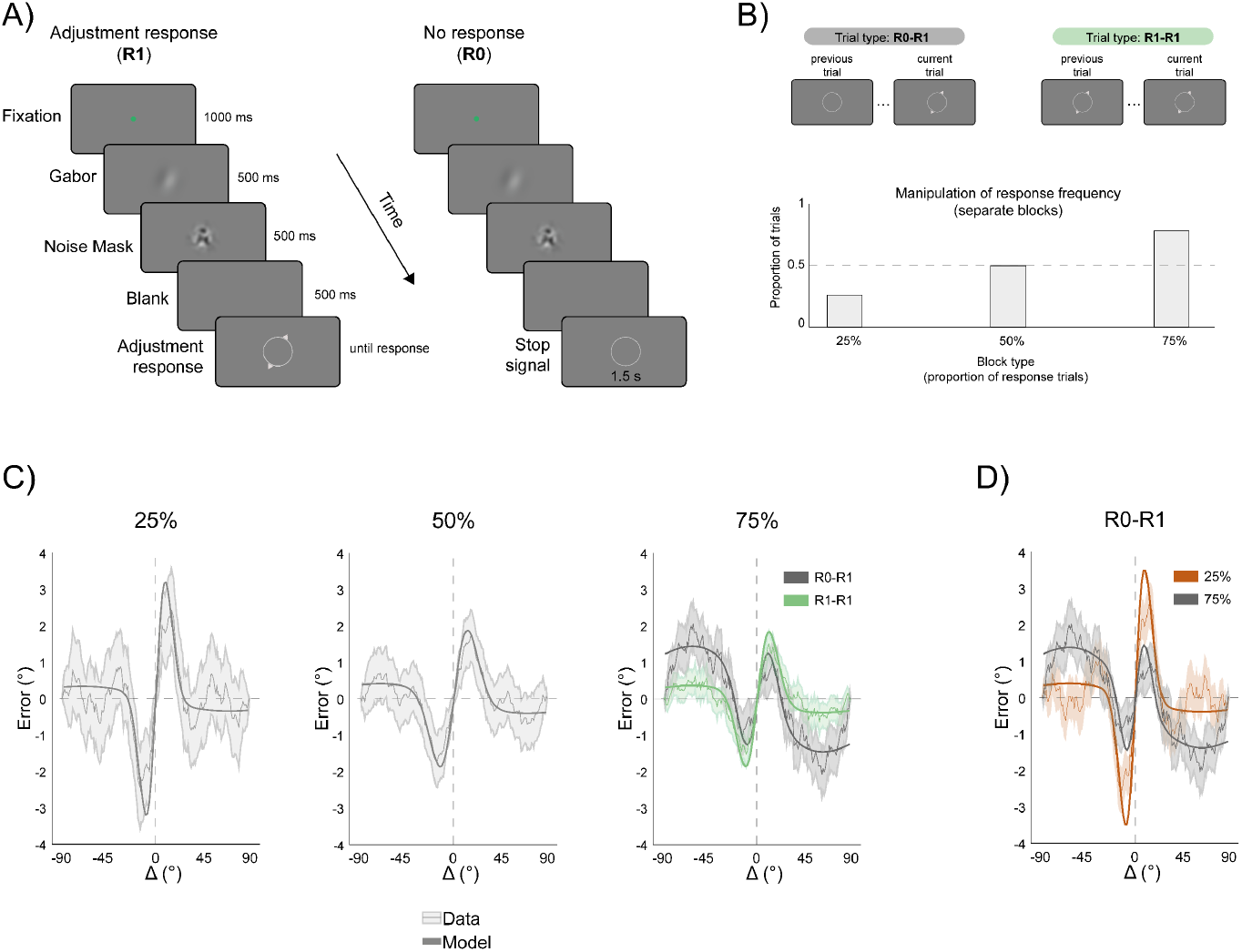
Experiment 1: Paradigm and behavioral results. A) Example of the sequence of events in the two trial types. In R1, an oriented Gabor was followed by a mask, and after a brief blank interval participants were presented with a response tool—a circular frame with two triangles on opposite ends—which they rotated to reproduce the perceived orientation. In R0, no triangles were shown on the response tool, and participants passively awaited the next trial, as no response was required. B) Main conditions of interest for measuring serial dependence. In R0–R1, serial dependence was measured from a previous trial that required no response; in R1–R1, it was measured between two consecutive response trials. Across separate blocks (bottom bar plot), we varied the ratio of response to no-response trials: 25% responses (rare response trials), 50% responses (equal proportion of response and no-response trials), and 75% responses (frequent responses, rare no-response trials). C) Serial dependence in orientation adjustment errors across blocks. Errors (y-axis) are plotted as a function of the orientation difference between the stimulus in the previous and current trial (Δ, x-axis). Data show the circular running average of aggregated single-trial errors across participants, with shaded areas indicating ±1 SD. Fits result from model comparisons of different formulations of the sum of two derivative-of-Gaussian (*δ*oG) functions, with or without an additional effect of trial type (R0–R1 vs. R1–R1; see Methods). In plots showing a single curve and fit (25% and 50% conditions), model comparison favored a simpler model without a trial-type effect. In contrast, for the 75% condition (two curves), the preferred model included a trial-type effect, indicating differences in serial dependence between R0–R1 (gray) and R1–R1 (green) trials. D) Comparison of serial dependence between conditions with approximately equal numbers of trials: R0–R1 under the 25% response condition (orange) and R0–R1 under the 75% response condition (gray). This contrast isolates the effect of response frequency on serial dependence (see Results for model comparison).

We focused on serial dependence in trials following a response (R1–R1) and those following a no-response trial (R0–R1; see Figure 1B) and assessed how these effects varied across response-frequency conditions. To this aim, we fitted each condition data with an extended derivative-of-Gaussian (*δ*oG) model, implemented as the sum of two *δ*oG functions (see Methods). This approach allows both attractive and repulsive components of serial dependence to be captured within the same function ^32^.

To test whether serial dependence was modulated by the previous trial type (response vs. no-response), we included a trial-type modulator on the amplitude and width parameters of the two *δ*oG components. We then compared this full model to a reduced model without the trial-type modulator using the Bayesian Information Criterion (BIC), computing the difference ΔBIC = BIC(reduced) – BIC(full) (see Methods). Positive ΔBIC values indicate evidence favoring the more complex model, whereas negative values favor the simpler one.

Figure 1C shows the best-fitting model results across conditions. Evidence favored the full model in the 75% response condition (ΔBIC_75%_ = +3.19), indicating an effect of the previous trial type on serial dependence. In contrast, the 25% and 50% conditions showed evidence favoring the simpler model (ΔBIC_25%_ = –12.5, ΔBIC_50%_ = –8.70).

In the 25% and 50% conditions serial dependence was predominantly attractive, exhibiting the typical positive bias toward the previous orientation. In the 75% condition, serial dependence was attractive following response trials (R1–R1). However, following no-response trials (R0–R1), the pattern showed stronger repulsive tails, consistent with prior findings ^18,33^.

Because the number of R0–R1 and R1–R1 trials differed across response-frequency conditions (e.g., R0–R1 trials were rarer in the 75% condition), we performed an additional comparison equating trial counts. Specifically, we compared R0–R1 trials from the 25% condition (≈ 18.2% trials) to R0–R1 trials from the 75% condition (≈ 18.1% trials). This confirmed a robust effect of overall response frequency (ΔBIC = +4.74), with stronger repulsive components observed when no-response trials were rare (75% condition).

Taken together, these behavioral results indicate that serial dependence is modulated by the presence or absence of a response on the preceding trial, and that this modulation depends on the global frequency of response trials. When no-response trials are rare, serial dependence following such trials shows reduced attraction and enhanced repulsion, particularly for larger Δ values.

### Experiment 2

What can explain the effect of previous response requirements and their dependence on the frequency of responses? One possibility is that after rare no-response trials, participants come to expect that a response will be required on the next trial. This interpretation aligns with extensive evidence showing that trial-by-trial expectations modulate behavior across perceptual and decision-making tasks ^31,34^. Such expectations may prompt a re-engagement with the task and enhanced processing of the current stimulus following a no-response event.

To further investigate this possibility, we analyzed an independent EEG dataset from a similar orientation adjustment task in which response and no-response trials occurred with equal probability (50%; see Methods and Figure 2A). Behaviorally, participants again showed modulation of serial dependence by previous trial type, with reduced attraction and increased repulsion after no-response trials (ΔBIC = +8640; Figure 2B), consistent with Experiment 1 despite the different response frequency conditions.

**Figure 2.**
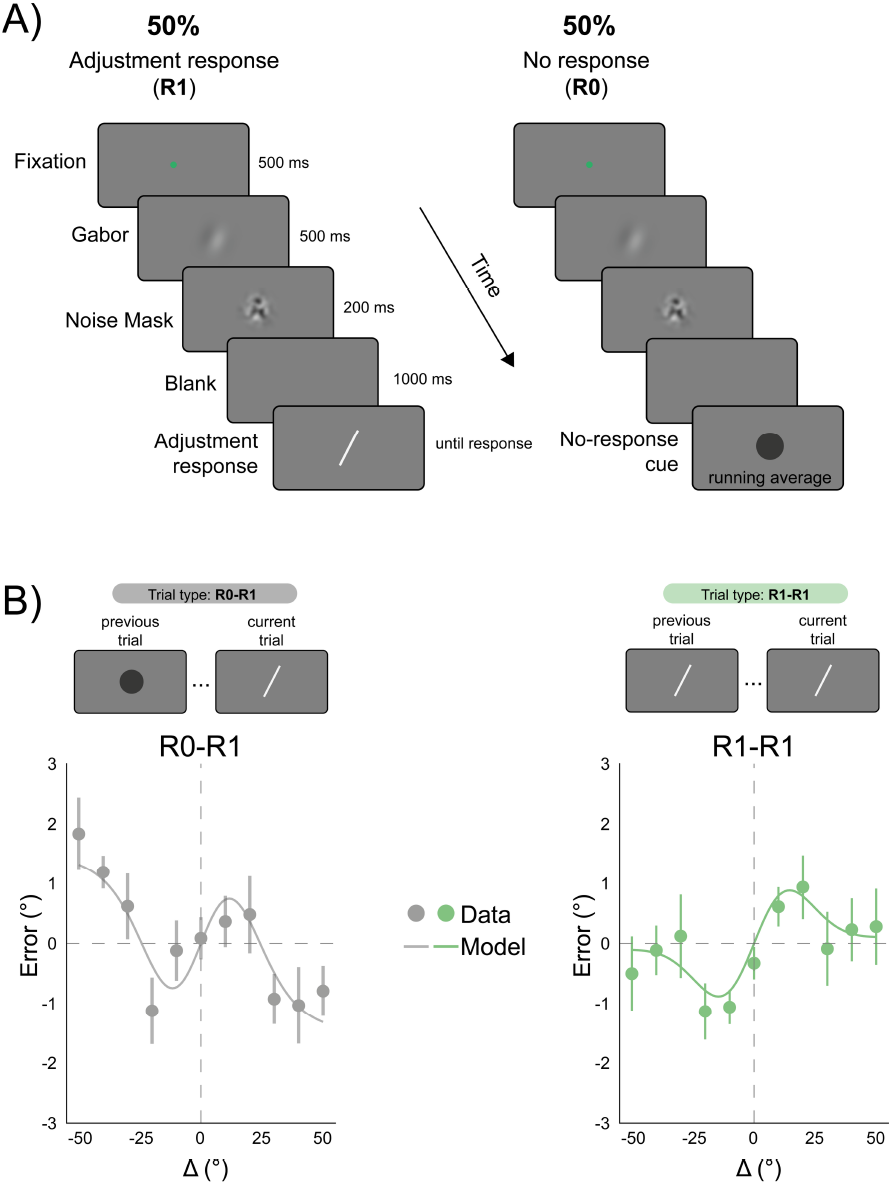
Experiment 2: Paradigm and behavioral results. A) Sequence of events in the two trial types. In this EEG experiment, the response frequency was 50%, meaning R0 and R1 trials were equally likely to occur. In R1 trials, participants reproduced the perceived orientation by rotating a response bar, whereas in R0 trials, a black disk was presented instead of the response tool. B) Serial dependence in orientation adjustment errors in the two types of trials (R0–R1, following no-response trials, in gray; R1–R1, following response trials, in green). Dots represent the mean error across participants for each Δ value (shown in discrete steps; see Methods). Error bars indicate SEM corrected for repeated measures. The curves show the predictions of the best-fitting model, following the same approach as in Experiment 1.

We next focused on the analysis of EEG activity time-locked to the current stimulus comparing the two trial types. Global Field Power (GFP, see Methods) was significantly higher following no-response trials (R0–R1) than following response trials (R1–R1) in a time window from 155 ms to 310 ms after stimulus onset (Figure 3A). This time range and the associated topography difference between conditions (Figure 3B) appear to be suggestive of relatively late attentional modulations (e.g., involving time windows characteristics of the P1/N1 complex). A linear model predicting trial-by-trial bias from GFP and trial type (R0–R1 vs. R1–R1, see Methods) revealed a significant positive effect of trial type, indicating larger positive (attractive) bias in orientation adjustment responses in the R1–R1 condition compared to R0–R1 (β = 0.83 ± 0.21 SE, *t*(5517) = 3.92, *p* <.001). In contrast, GFP fluctuations were negatively associated with bias magnitude (β = – 0.85 ± 0.34 SE, *t*(5517) = –2.48, *p* =.013), suggesting that higher GFP was predictive of stronger repulsive biases (Figure 3C). The overall model fit was modest (R^2^ = 0.004, F(2, 5517) = 11.1, *p* <.001).

**Figure 3.**
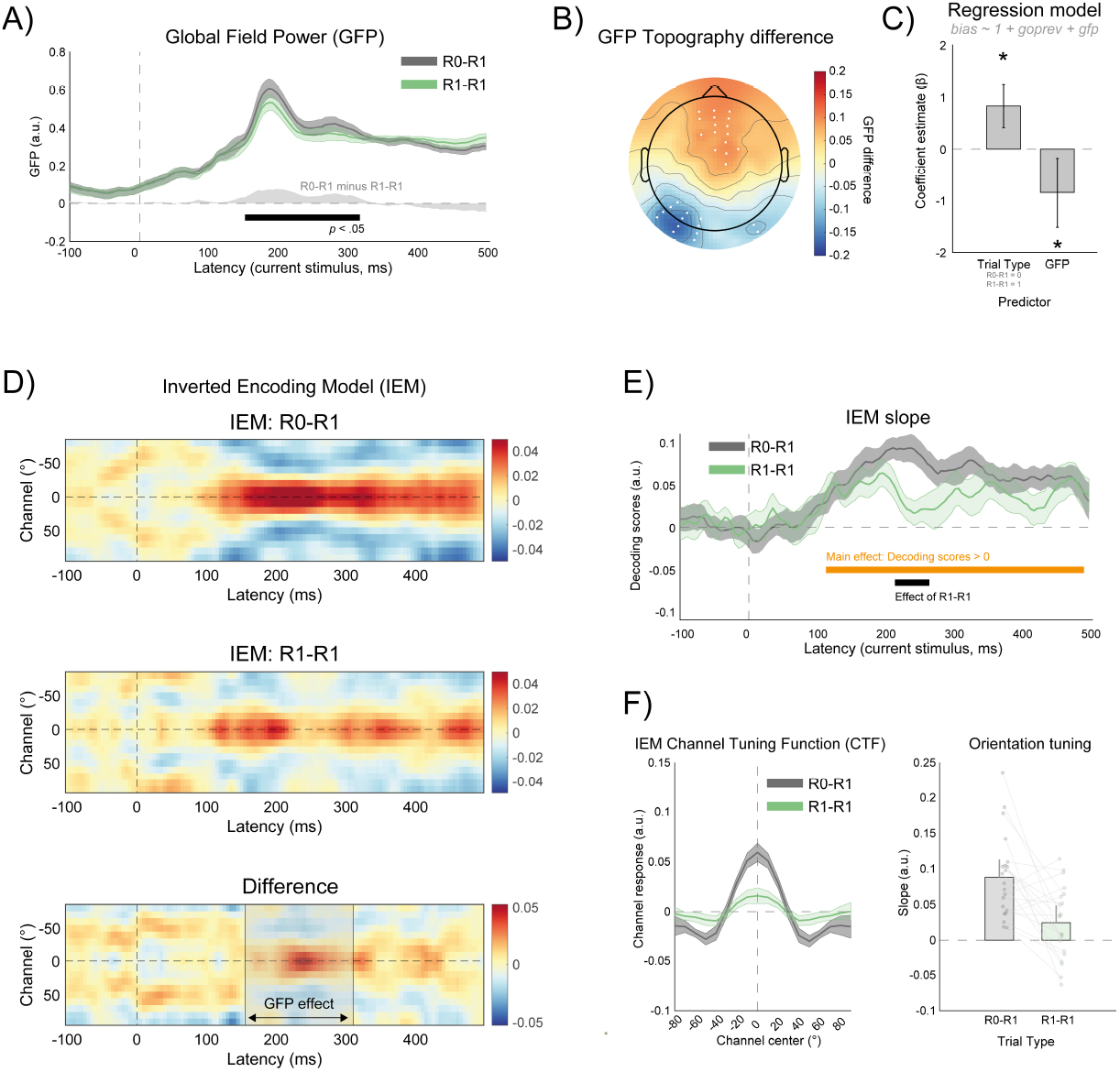
Experiment 2: EEG results. A) Global Field Power (GFP) time-locked to the current stimulus by trial type, averaged across participants. GFP was higher in the R0–R1 condition (gray) compared to the R1–R1 condition (green) within a time window from 155 to 310 ms. Shaded regions indicate SEM corrected for repeated measures. The light gray trace shows the GFP difference between conditions, with the significant time window marked by a black horizontal line (see Results). B) Scalp topography of the GFP difference between trial types, with electrodes showing peak positive or negative differences (top 25% percentile) highlighted by white circles. C) Linear model coefficients (error bars are the 95% CI) predicting bias in adjustment errors (positive = attraction toward the previous stimulus; negative = repulsion) as a function of trial type and trial-by-trial GFP fluctuations within the significant time window identified in (A). Asterisks denote significant coefficients (p <.05). D) Inverted Encoding Model (IEM) results showing reconstructed channel responses over time (x = time relative to stimulus onset; y = IEM channels), separately for R0–R1 and R1–R1 trials, and their difference (bottom row). The gray shaded rectangle indicates the GFP difference window reported in (A). E) Decoding scores (derived from the slope of the channel tuning function; see Methods) for the two trial types, using the same color coding as in (A). Decoding of the current orientation was significant from 125 ms onward (orange line) and differed between trial types in a narrower window from 215 to 260 ms (black line). Shaded regions are corrected SEM. F) During the time window showing a significant trial-type effect on decoding, channel tuning functions (CTFs) were amplified in the R0–R1 condition (left; shaded regions = corrected SEM), indicating steeper slopes and enhanced orientation representations (right; dots represent individual participants; error bars show 95% CI corrected for repeated measures).

Finally, we assessed how the current trial orientation was represented in EEG activity using an inverted encoding model (IEM; see Methods). Orientation decoding was significant from 120 ms to 485 ms post-stimulus (Figure 3D-E). Importantly, decoding accuracy was higher for R0–R1 than R1–R1 trials in the time window from 215 ms to 260 ms (Figure 3E), accompanied by increased amplitude of the reconstructed channel response functions (Figure 3F), indicating stronger orientation-selective responses following no-response trials. Together, these results suggest that after no-response trials, EEG activity and the strength of orientation representations are enhanced.

## Discussion

In two experiments, we investigated how trial-by-trial response expectations influence serial dependence and EEG activity. In a behavioral study (Experiment 1), we manipulated the proportion of response trials in separate blocks and found that serial dependence was modulated primarily after rare no-response trials (i.e., in the 75% response block). In the EEG study (Experiment 2), we measured changes in EEG signal strength and orientation decoding in the current trial, depending on the previous trial response requirement. Again, we observed a modulation of serial dependence in behavior, accompanied by distinct changes in EEG activity and its orientation-selective content. Together, these results suggest that no-response trials trigger a re-engagement with the task and enhanced processing of the current stimulus.

In Experiment 1, the effect was evident in the condition where no-response trials were rare (the 75% response condition), manifesting as a reduction in attractive serial dependence and an increase in repulsive bias after no-response trials. The attractive and repulsive components of serial dependence have previously been linked to effects of prior history at different processing stages: repulsion has been linked to adaptation-like aftereffects at sensory levels, whereas attraction has been related to the persistence of representations at higher levels such as decision-making or working memory ^18,25,26,35,36^. Within this framework, the dominance of one component over the other would depend on factors such as the strength of the previous stimulus (e.g., duration, contrast) and the extent to which it was attended, maintained in memory, or reported.

Although this hierarchical distinction remains debated ^see 3 for a review^, our results are broadly consistent with the idea that prior responses modulate the balance between these opposing forces. One possibility is that withholding a response on the previous trial, and the explicit cue indicating that no response is required, leads to a “reset” or active removal of the most recent stimulus trace ^15,37^. In a hierarchical view, this would reduce the influence of prior events at higher-level processing stages (attractive biases) and consequently allow low-level adaptation to dominate, producing repulsive effects ^15,18,24^. Alternatively, active removal of no longer relevant information from working memory, as well as stimulus devaluation processes following response withholding, could themselves induce repulsive biases without invoking sensory adaptation mechanisms ^15,18,37^.

However, these interpretations alone cannot explain why the effects of previous trial type, at least in Experiment 1, emerged only when no-response trials were rare. Consistent with our hypothesis, we propose that these effects reflect expectations to respond on the current trial following a no-response trial. When response trials are frequent, such expectations are stronger, leading to a re-engagement with the task and current stimulus. This, in turn, enhances stimulus processing, reduces the influence of prior events, and increases the tendency to discriminate the current stimulus from previous ones—manifesting as stronger repulsive biases. One plausible mechanism underlying this effect is a refocusing of attention toward the current sensory input. This interpretation aligns with evidence that attractive serial dependence tends to occur under conditions of reduced sensory processing or higher uncertainty, for instance when stimuli are weak, brief, or task resources are limited ^6,15,38^. Our findings demonstrate that such modulations can arise entirely from internal states of the observer, driven by expectations about upcoming task demands, even when the physical properties of the stimuli are constant ^see also 39,40^.

In Experiment 2, the EEG results provided a further, complementary characterization of this phenomenon. We observed increased overall EEG signal strength and higher orientation decoding scores for the current stimulus following no-response trials. These findings, evident in the GFP and IEM analyses respectively, indicate that orientation-selective responses were amplified after no-response trials. The time window of the GFP effect (155-310 ms post-stimulus) and its scalp distribution correspond to typical EEG components modulated by attention, including the P1 and visual N1. Likewise, the IEM results within this period revealed increased amplitude of the reconstructed channel response functions after no-response trials, consistent with attentional gain modulation ^41^. Together, these results support the interpretation that following no-response trials, enhanced attentional engagement and increased allocation of processing resources strengthen the neural representation of the current stimulus.

It is worth noting that, behaviorally, these effects emerged under different conditions in the two experiments: when no-response trials were rare (Experiment 1, 75% response) and when response and no-response trials were equally likely (Experiment 2, 50%). Although the precise nature of this difference remains to be determined, the consistency of the pattern suggests a robust phenomenon ^see also Figure 3 in 18^. It is possible that broader contextual factors, such as the overall structure of the experiment (multiple response-ratio blocks in Experiment 1 vs. a fixed 50% design in Experiment 2), might have contributed to participants’ expectations, pointing to an additional higher-level contextual modulation of serial dependence by experimental design itself.

We propose that expectations naturally arising in contexts with intermixed response and no-response trials may be what induce these fluctuations in attention and task engagement, which in turn modulate the balance between attractive and repulsive serial dependence. This perspective departs from most prior studies, which have treated previous responses either as motor confounds ^1,27^ or as key contributing factors in serial dependence ^18,42^, without identifying the processes responsible for trial-by-trial modulations.

A central implication is that attractive serial dependence—in some frameworks interpreted as evidence for a dedicated mechanism stabilizing perception ^1,43^—does not result from a fixed and hard-wired process. Rather, it is shaped by top-down factors such as dynamic fluctuations in task engagement and expectation, beyond a fixed mechanism. In this sense, our findings support the view that attractive serial dependence emerges under specific internal states that favor the persistence and interference of previous task content with current decisions ^39,40^. These states alternate with states that place greater weight on current sensory input. Our manipulations may have specifically targeted this transition: from a state promoting attraction to prior stimuli (during continuous responding) to one enhancing current-stimulus processing and reducing interference from prior events (following a no-response trial, when a response is expected next). To our knowledge, we provide the first evidence of neural correlates associated with this putative state transition, marked by increased EEG activity and amplified orientation tuning for the current stimulus. Previous decoding studies have primarily focused on decoding residual traces of past stimuli or the bias they induce in current neural representations ^22,26,35,44^, but none have reported changes in current-stimulus representations linked to trial-by-trial fluctuations in serial dependence. The demonstration that serial dependence is dynamically modulated by internal states—with identifiable EEG signatures of these fluctuations— offers a novel framework for understanding inter-individual variability in serial dependence, as well as deviations observed in clinical populations ^28,45,46^.

In sum, we show that trial-by-trial response expectations represent a key source of modulation in serial dependence, with no-response trials triggering re-engagement with the task and enhanced processing of the subsequent stimulus, ultimately reweighting the balance between attractive and repulsive influences from recent history.

## Methods

### Experiment 1

#### Participants

The study was conducted at the École polytechnique fédérale de Lausanne (EPFL, Lausanne, Switzerland). Twenty-four healthy participants (11 females, age range: 21-37 y) were recruited for monetary reward (20 CHF/hour). All participants had normal or corrected-to-normal vision according to the Freiburg Visual Acuity test ^47^ and received written informed consent before the experiment. The study was approved by the local ethics committee in accordance with the Declaration of Helsinki (World Medical Organization, 2013).

#### Apparatus

All experiments were run on a gamma-corrected VG248QE monitor (resolution: 1920 x 1080 pixels, refresh rate: 120 Hz) in a darkened room. Stimuli were generated with custom-made scripts written in MATLAB (R2013a) and the Psychophysics Toolbox and presented on a grey background (62.66 cd/m^2^). Participants sat at 57 cm from the computer screen, with their head on a chin rest.

#### Stimuli and procedure

Figure 1A depicts the sequence of events in an experimental trial. Each trial started with a fixation spot shown for 1000 ms. After fixation, a low contrast (peak contrast of 25% Michelson) and low spatial frequency (0.33 cycles per degree) Gabor (Gaussian envelope: 1.5°) was shown for 500 ms, followed by a high-contrast noise mask (95% Michelson) for other 500 ms. The Gabor could have any orientation in the 0-179° circular space, randomly assigned on each trial. A blank interval (500 ms) preceded the appearance of the response tool or the stop signal. The response tool was made of a gray circular frame (diameter of 4°) with two triangles (e.g., arrowheads) positioned on the outline, at two equidistant extremities of an imaginary line. In stop trials, the triangles were not shown, and the circular frame alone indicated the absence of a response.

In response trials, participants had to rotate the orientation of the imaginary line to match the perceived orientation of the Gabor, by moving the mouse in the upward and downward directions, confirming the response with a left click. In stop trials, they were instructed to withhold the response and wait until the next trial. The waiting time before the next trial was calibrated to the running average of individual adjustment times.

The main manipulation consisted in varying the proportion of response and stop trials in three separate blocks. As shown in Figure 1C, the frequency of responding was either 25%, 50%, or 75%. The order of blocks was counterbalanced across participants.

Before the experiment, all participants were provided written and verbal instructions. To ensure that participants understood the task, they performed a brief practice before the experiment, under the supervision of the experimenter. The experiment consisted of 3 blocks, for a total of 600 trials, lasting approximately 1 hour.

#### Behavioral data analysis

The analysis of behavioral serial dependence focused on adjustment errors, defined as the acute angular difference between the reported orientation and the stimulus orientation on each trial. Prior to the analysis, errors were cleaned through a multi-step procedure.

First, trials with absolute errors larger than 45° were classified as lapses and excluded ^28^. Trials with adjustment response times longer than 10 seconds were also excluded. Second, the remaining errors were demeaned and corrected for systematic orientation-dependent biases using a sinusoidal residualization approach ^18,48^. Specifically, a sum of six sine and cosine functions was fitted to the error as a function of the current orientation using the MATLAB fit.m function with the model specification “sin6”, and the resulting fit was subtracted from the errors. Next, errors identified as statistical outliers—defined as values more than 1.5 interquartile ranges above the upper quartile or below the lower quartile—were removed using the isoutlier function in MATLAB with the “quartiles” method. Across all steps, less than 5% of response trials were excluded from the analysis. The remaining cleaned data were used to assess serial dependence effects.

To quantify serial dependence and compare its pattern across conditions, we modeled the relationship between current-trial errors and Δ (the relative orientation between the current and previous stimuli), using a parametric model based on the first derivative of a Gaussian (*δ*oG) ^1^:

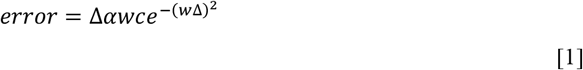

where 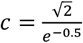 is a normalization constant, *w* controls the inverse width of the curve, and α is the amplitude parameter indicating the strength and direction of the bias. Positive values of α reflect an attractive bias (i.e., errors shifted toward the previous stimulus), while negative values indicate a repulsive bias (i.e., errors shifted away).

Previous studies have shown that serial dependence can exhibit both attractive and repulsive components, with attraction typically occurring for small Δ values and repulsion for larger Δs ^32,33^. Moreover, manipulations of prior response requirements have revealed increased repulsion at large Δ following non-response (R0) trials ^see Experiment 3 in 18^. Because a single *δ*oG cannot capture both attractive and repulsive components simultaneously, we first visually inspected whether the data, overall across conditions, displayed a biphasic pattern consistent with both effects. After confirming this pattern, a “double *δ*oG” model—comprising the weighted sum of two *δ*oG functions with free amplitude and width parameters—was fit to the error data ^32^.

To assess whether the shape and strength of serial dependence varied with trial history, we implemented a condition-dependent model including interaction terms between Δ and the response condition (R0-R1 vs. R1-R1). This allowed both the amplitude and width parameters of the two *δ*oG components to vary as a function of condition.

Model parameters were estimated using constrained nonlinear least squares optimization (fmincon in MATLAB) applied to the pooled single-trial data across participants. Model comparison was performed using the Bayesian Information Criterion (BIC), which penalizes model complexity. Differences in BIC (ΔBIC) greater than 2 are considered positive evidence against the model with the higher BIC ^49^. We specifically tested whether a model allowing different parameters across conditions provided a better fit, indicating that the presence or absence of a response on the preceding trial significantly altered the serial dependence pattern observed on the current trial.

### Experiment 2

#### Participants

Experiment 2 was conducted at the University of Fribourg (Fribourg, Switzerland). Twenty-one healthy participants (17 females, age range: 18-31 y) were recruited from the local student population. All participants had normal or corrected-to-normal vision, verified prior to the experiment using the Freiburg Acuity Test, for which a minimum value of 1 was required with both eyes open ^47^. Written informed consent was obtained from all participants prior to participation. The experimental procedures were approved by the regional ethics board (CER-VD, Protocol Nr. 2016-00060) and were conducted in accordance with the Declaration of Helsinki.

#### Apparatus

Participants were seated in a dark, electrically shielded room, at a distance of 80 cm from the display monitor. EEG was recorded using a 128-channel BioSemi ActiveTwo system (BioSemi, Amsterdam, The Netherlands). Stimuli were generated using MATLAB (MathWorks, Natick, MA) and the Psychophysics Toolbox ^50^ and presented on a VIEWPixx/3D display system (1920 × 1080 pixels resolution; 120 Hz refresh rate; VPixx Technologies, Canada).

#### Stimuli and procedure

An example trial sequence and the two main conditions are illustrated in Figure 2A. Each trial began with a fixation point displayed at the center of the screen for 500 ms. This was followed by a Gabor patch presented for 500 ms, which was then masked by a noise pattern lasting 200 ms. A blank screen was shown for 1000 ms before the response stage.

In R1 trials, participants were shown a randomly oriented white bar and asked to rotate it to match the perceived orientation of the preceding Gabor stimulus using the left and right arrow keys on a keyboard. The final response was confirmed by pressing the spacebar. In R0 trials (50% of trials, randomly assigned), a central gray oval appeared in place of the response tool, serving as an explicit no-response cue. Participants were instructed to await the next trial without providing any response.

The duration of the no-response cue was initially fixed at 1.5 s, based on average adjustment durations reported in similar tasks ^1,18^, but was subsequently adapted after the first 50 trials to reflect the running average of the participant’s actual adjustment times. The inter-trial interval (blank screen) was randomly jittered between 1.1 and 1.3 s in 20 ms increments.

Gabor stimuli were sinusoidal gratings (spatial frequency: 1 cycle/°; Michelson contrast: 0.25) windowed by a Gaussian envelope with a sigma of 1°. Stimuli were presented centrally on a gray background. The mask consisted of full-contrast Gaussian-filtered white noise. On each trial, the orientation of the Gabor was selected at random from 0° to 170° in 10° increments, with the constraint that the orientation difference relative to the preceding trial (Δ: previous minus current orientation) did not exceed ±50°. This constraint resulted in 11 possible Δ values ranging from –50° to +50° in 10° steps.

#### Behavioral data analysis

The analysis of behavioral serial dependence followed the procedure and steps described for Experiment 1. Across all steps, less than 6% of response trials were excluded from the analysis.

#### EEG recording and preprocessing

EEG data were recorded at a sampling rate of 2048 Hz. Signal quality was monitored online by ensuring that offsets between active electrodes and the Common Mode Sense – Driven Right Leg (CMS–DRL) feedback loop remained within ±20 mV.

Offline preprocessing was conducted in EEGLAB 2019.1 ^51^. EEG data were first downsampled to 200 Hz using pop_resample.m, with an anti-aliasing filter cutoff of 0.8 and a transition bandwidth of 0.4 (normalized units in π rad/sample). Data were locally detrended using the PREP pipeline plugin ^52^. Line noise at 50 Hz and its harmonics was removed using pop_cleanline.m.

Data were then epoched from -1000 to 2000 ms relative to stimulus onset. Bad channels and epochs were identified by visual inspection and excluded prior to further preprocessing.

Remaining physiological artifacts were identified using independent component analysis (ICA) with the Infomax algorithm as implemented in EEGLAB’s runica function. To mitigate rank deficiency and reduce computational load, the data were first reduced in dimensionality using principal component analysis (PCA), retaining components that explained more than 98% of the variance.

Artifact components were classified using the MARA (Multiple Artifact Rejection Algorithm), and those explaining more than 90% of the total variance were inspected visually and removed. Subsequently, bad channels were interpolated using the nearest-neighbor spline method, and data were re-referenced to the average reference. On average, 49.76±33.19 (7.54%) epochs were marked as outliers and excluded, 20.14±7.66 (15.74%) channels interpolated, and 14.43±6.72 (13.33%) ICA removed.

#### EEG analysis

##### Global Field Power (GFP)

To compare EEG evoked activity as a function of the previous trial condition (R0–R1 vs. R1–R1), we computed the Global Field Power (GFP) across electrodes. GFP is a reference-independent measure of the overall strength of scalp EEG activity at each time point, defined as the standard deviation of the voltage potentials across all electrodes ^53^. The analysis was restricted to the time window from -100 to 500 ms after stimulus onset.

Here, GFP time courses were computed separately for each participant and condition, following baseline normalization using the 100 ms pre-stimulus interval. This normalization step controlled for potential differences in overall scalp amplitude due to residual motor-related activity following R1–R1 trials. Only behaviorally valid trials were included in the analysis. In addition to GFP, single-trial EEG data and channel-wise ERP maps were extracted for subsequent analyses.

To test for time-resolved differences in GFP between trial types, we employed non-parametric cluster-based permutation testing with 1000 sign-flip permutations (flipping the GFP difference sign within each participant). Cluster-level statistics—computed as the maximum summed *t*-values across temporally contiguous points—were compared to a null distribution to determine significance, providing intrinsic correction for multiple comparisons ^54^. Temporal clusters with *p* <.05 (two-tailed) were considered statistically significant.

To identify the topographic distribution of GFP differences, we computed the average scalp map of the R0–R1 vs. R1–R1 GFP difference, averaged over the time window corresponding to the significant temporal cluster.

##### GFP-Behavior relationship

To assess whether trial-by-trial fluctuations in global field power (GFP) predicted behavioral biases, we extracted single-trial GFP values within the significant time window of the effect (by averaging over time and computing the standard deviation across electrodes for each trial). As a measure of behavioral bias, we defined an individual-level global bias index as the product of signed error and Δ (i.e., error × sign of orientation difference), excluding trials where Δ = 0°. This index—akin to the error-folding procedures employed in previous studies ^8^—captures the direction and magnitude of systematic deviations in response errors toward (positive bias) or away from (negative bias) the previous-trial orientation. It thus provides a single-trial measure of serial dependence for each participant and condition.

We modeled the relationship between GFP and the bias index using a linear regression predicting trial-by-trial bias from GFP, including an additional predictor coding for the condition (R0–R1 vs. R1–R1, coded as a binary variable: 0–1) and an intercept. This model allowed us to estimate regression coefficients quantifying the relationship between GFP fluctuations and the magnitude of serial dependence, while accounting for differences between experimental conditions.

##### Inverted Encoding Model (IEM)

To assess whether the quality of orientation representations in EEG activity was modulated by trial history (e.g., previous response condition), we applied an Inverted Encoding Model ^IEM; 55,56^ from -100 to 500ms.

The IEM assumes that stimulus orientation is encoded by a set of hypothetical neural channels with idealized tuning functions. We used 18 orientation channels, each spaced 10° apart, covering orientations from 0° to 170°. Each channel’s tuning curve was modeled as a half-wave rectified cosine raised to the 18th power.

For each trial, we generated a predicted pattern of channel responses based on the presented stimulus orientation, forming a channel response matrix *C* of size [k × n] (with *k* = 18 channels and *n* = number of trials). The observed EEG data were modeled as:

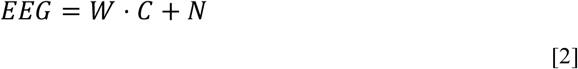

where *EEG* is the electrode × trial matrix [m x n] (with *m* = 128 electrodes), *W* is the weight matrix [m x k] mapping channel responses to scalp sensors, and *N* is residual noise. Weights were estimated using least-squares regression on a training set:

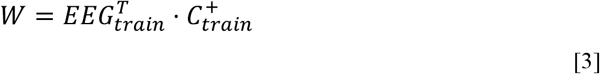

where 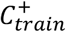 is the pseudoinverse of the training channel response matrix. Orientation-selective channel responses for the test data were reconstructed as:

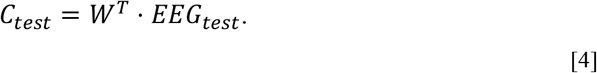

The IEM was applied using a sliding window of 10 samples (corresponding to 50 ms), using a 3-fold cross-validation procedure with 300 iterations per subject and condition. The decoder was trained separately for each experimental condition (R0-R1 and R1-R1).

For each time point, the reconstructed channel tuning functions (CTFs) were circularly realigned so that the channel tuned to the presented stimulus orientation was centered at 0°. This yielded a [time x channel] matrix per subject.

Decoding performance was quantified as the slope of a linear fit between the reconstructed CTF and an idealized basis function centered at 0°. This decoding score (CTF slope) reflects how well the reconstructed signal matches the expected orientation tuning. Trials excluded from the behavioral analysis (e.g., outliers in *Behavioral data analysis*) were also excluded from the IEM analysis.

##### IEM surrogate distribution and null hypothesis testing

To assess the statistical significance of decoding performance, we constructed a surrogate distribution of CTF slopes under the null hypothesis of no orientation-specific information. Surrogates were generated by shuffling the orientation labels during the circular realignment step, thereby disrupting the systematic alignment between reconstructed CTFs and true orientations. This procedure was repeated 300 times per subject, yielding a null distribution of 300 surrogate slopes per time point.

To test whether decoding performance (CTF slope) differed from zero and whether it varied by condition (R0-R1 vs. R1-R1), we performed a regression analysis at each time point. For each subject, CTF slope values across both conditions were concatenated into a single response vector. A binary predictor variable coded the experimental condition (0 for R0-R1 and 1 for R1-R1). Regression coefficients (intercept and condition effect) and their associated t-statistic were estimated using least-squares regression.

To evaluate the group-level significance of these effects, we applied the same regression procedure to the surrogate slope distributions, producing a null distribution of t-statistics associated with the regression coefficients (intercept and condition) per time point. To correct for multiple comparisons across time points, we used a cluster-based permutation approach. For each surrogate iteration, we identified time points where the empirical t-statistics for the intercept and condition effect exceeded the two-tailed critical value (|*t*| > *t*_α/2_, with *α* = 0.05 and degrees of freedom *df* = 20). We then computed the sum of supra-threshold t-values within each cluster to derive a cluster-level statistic and retained the maximum absolute cluster sum from each iteration to form a null distribution. Cluster-level statistics from the real data were calculated using the same procedure. P-values for observed clusters were obtained by comparing these statistics to the corresponding null distributions for positive and negative clusters. Significant clusters in the intercept term identified periods during which the current orientation was reliably decoded from EEG activity. A significant condition effect indicated temporal windows where the quality of orientation representations differed between the R0-R1 and R1-R1 conditions. Specifically, a negative condition coefficient reflected reduced decoding fidelity in the R1-R1 condition relative to R0-R1.

## Acknowledgments

The authors thank Gizay Ceylan for assistance with the setup and data collection of Experiment 1, Laura Cohen for data collection, and Laura Cohen and Cemre Yilmaz for the initial preprocessing and data handling of Experiment 2. We also thank Gijs Plomp for providing the resources to run Experiment 2.

## Funding

Data for Experiment 2 were collected while D. Pascucci was a postdoctoral researcher in the lab of G. Plomp (University of Fribourg) with support from the Swiss National Science Foundation (grants PZ00P3_131731 and PP00P1_157420). Data collection for Experiment 1 and the final analyses were supported by the Swiss National Science Foundation (grants PZ00P1_179988, PZ00P1_179988/2, and TMSGI1_218247).

## Author contributions

Experiment design: D.P.

Data analysis: D.P., J.L

Writing manuscript: D.P., J.L

Illustrations: D.P.

Editing manuscript: D.P., J.L

## Competing interests

The authors have no conflicts of interest to declare.

## Data availability

Data and code will be made available before publication.

## Notes

### Competing Interest Statement

The authors have declared no competing interest.

## References

1. Fischer, J. & Whitney, D. Serial dependence in visual perception. Nat. Neurosci. 17, 738–743 (2014).

2. Manassi, M., Murai, Y. & Whitney, D. Serial dependence in visual perception: A meta-analysis and review. J. Vis. 23, 18 (2023).

3. Pascucci, D. et al. Serial dependence in visual perception: A review. J. Vis. 23, 9 (2023).

4. Ceylan, G., Herzog, M. H. & Pascucci, D. Serial dependence does not originate from low-level visual processing. Cognition 212, 104709 (2021).

5. Cicchini, G. M., Mikellidou, K. & Burr, D. The functional role of serial dependence. Proc. R. Soc. B 285, 20181722 (2018).

6. Gallagher, G. K. & Benton, C. P. Stimulus uncertainty predicts serial dependence in orientation judgements. J. Vis. 22, 6–6 (2022).

7. Tanrikulu, Ö.D., Pascucci, D. & Kristjánsson, Á. Stronger serial dependence in the depth plane than the fronto-parallel plane between realistic objects: Evidence from virtual reality. J. Vis. 23, 20 (2023).

8. Barbosa, J. & Compte, A. Build-up of serial dependence in color working memory. Sci. Rep. 10, 10959 (2020).

9. Fischer, C. et al. Context information supports serial dependence of multiple visual objects across memory episodes. Nat. Commun. 11, 1–11 (2020).

10. Kim, S., Burr, D. & Alais, D. Attraction to the recent past in aesthetic judgments: A positive serial dependence for rating artwork. J. Vis. 19, 19–19 (2019).

11. Liberman, A. & Whitney, D. The serial dependence of perceived emotional expression. J. Vis. 15, 929–929 (2015).

12. Kiyonaga, A., Scimeca, J. M., Bliss, D. P. & Whitney, D. Serial Dependence across Perception, Attention, and Memory. Trends Cogn. Sci. (2017).

13. Lõoke, M., Guérineau, C., Broseghini, A., Mongillo, P. & Marinelli, L. Visual continuum in non-human animals: serial dependence revealed in dogs. Proc. R. Soc. B Biol. Sci. 291, 20240051 (2024).

14. Papadimitriou, C., Ferdoash, A. & Snyder, L. H. Ghosts in the machine: memory interference from the previous trial. J. Neurophysiol. 113, 567–577 (2014).

15. Ceylan, G. & Pascucci, D. Attractive and repulsive serial dependence: The role of task relevance, the passage of time, and the number of stimuli. J. Vis. 23, 8 (2023).

16. Guan, S. & Goettker, A. Individual differences reveal similarities in serial dependence effects across perceptual tasks, but not to oculomotor tasks. J. Vis. 24, 2–2 (2024).

17. Kim, S. & Alais, D. Individual differences in serial dependence manifest when sensory uncertainty is high. Vision Res. 188, 274–282 (2021).

18. Pascucci, D. et al. Laws of concatenated perception: Vision goes for novelty, decisions for perseverance. PLOS Biol. 17, e3000144 (2019).

19. Alais, D., Leung, J. & Van der Burg, E. Linear summation of repulsive and attractive serial dependencies: orientation and motion dependencies sum in motion perception. J. Neurosci. 37, 4381–4390 (2017).

20. Blondé, P., Kristjánsson, Á. & Pascucci, D. Tuning perception and decisions to temporal context. iScience 108008 (2023) doi:10.1016/j.isci.2023.108008.

21. Fornaciai, M. & Park, J. Spontaneous repulsive adaptation in the absence of attractive serial dependence. J. Vis. 19, 21–21 (2019).

22. Luo, M., Zhang, H., Fang, F. & Luo, H. Reactivation of previous decisions repulsively biases sensory encoding but attractively biases decision-making. PLOS Biol. 23, e3003150 (2025).

23. Moon, J. & Kwon, O.-S. Attractive and repulsive effects of sensory history concurrently shape visual perception. BMC Biol. 20, 247 (2022).

24. Pascucci, D. & Plomp, G. Serial dependence and representational momentum in single-trial perceptual decisions. Sci. Rep. 11, 9910 (2021).

25. Fritsche, M., Spaak, E. & de Lange, F. P. A Bayesian and efficient observer model explains concurrent attractive and repulsive history biases in visual perception. eLife 9, e55389 (2020).

26. Sheehan, T. C. & Serences, J. T. Attractive serial dependence overcomes repulsive neuronal adaptation. PLoS Biol. 20, e3001711 (2022).

27. Manassi, M., Liberman, A., Kosovicheva, A., Zhang, K. & Whitney, D. Serial dependence in position occurs at the time of perception. Psychon. Bull. Rev. 1–9 (2018).

28. Pascucci, D. et al. Intact Serial Dependence in Schizophrenia: Evidence from an Orientation Adjustment Task. Schizophr. Bull. sbae106 (2024).

29. Cicchini, G. M., Mikellidou, K. & Burr, D. Serial dependencies act directly on perception. J. Vis. 17, 6–6 (2017).

30. Murai, Y. & Whitney, D. Serial dependence revealed in history-dependent perceptual templates. Curr. Biol. (2021).

31. Pascucci, D., Mastropasqua, T. & Turatto, M. Permeability of priming of pop out to expectations. J. Vis. 12, 21–21 (2012).

32. Houborg, C., Pascucci, D., Tanrikulu, Ö.D. & Kristjánsson, Á. The effects of visual distractors on serial dependence. J. Vis. 23, 1 (2023).

33. Fritsche, M. & de Lange, F. P. The role of feature-based attention in visual serial dependence. J. Vis. 19, 21 (2019).

34. Croson, R. & Sundali, J. The gambler’s fallacy and the hot hand: Empirical data from casinos. J. Risk Uncertain. 30, 195–209 (2005).

35. Fischer, C., Kaiser, J. & Bledowski, C. A direct neural signature of serial dependence in working memory. eLife 13, (2024).

36. Fritsche, M., Mostert, P. & de Lange, F. P. Opposite effects of recent history on perception and decision. Curr. Biol. 27, 590–595 (2017).

37. Shan, J. & Postle, B. R. The Influence of Active Removal from Working Memory on Serial Dependence. 5, 31 (2022).

38. Markov, Y. A., Tiurina, N. A. & Pascucci, D. Serial dependence: A matter of memory load. Heliyon 10, e33977 (2024).

39. Ozkirli, A. & Pascucci, D. It’s not the spoon that bends: Internal states of the observer determine serial dependence. 2023.10.19.563128 Preprint at 10.1101/2023.10.19.563128 (2024).

40. Weilnhammer, V., Murai, Y. & Whitney, D. Dynamic predictive templates in perception. Curr. Biol. 34, 4301–4306 (2024).

41. Reynolds, J. H. & Chelazzi, L. ATTENTIONAL MODULATION OF VISUAL PROCESSING. Annu. Rev. Neurosci. 27, 611–647 (2004).

42. Sadil, P., Cowell, R. A. & Huber, D. E. The push–pull of serial dependence effects: Attraction to the prior response and repulsion from the prior stimulus. Psychon. Bull. Rev. 1–15 (2023).

43. Collins, T. The perceptual continuity field is retinotopic. Sci. Rep. 9, 1–6 (2019).

44. Bae, G.-Y. & Luck, S. J. Reactivation of previous experiences in a working memory task. Psychol. Sci. 30, 587–595 (2019).

45. Barbosa, J. et al. Interplay between persistent activity and activity-silent dynamics in the prefrontal cortex underlies serial biases in working memory. Nat. Neurosci. 23, 1016–1024 (2020).

46. Bansal, S. et al. Qualitatively Different Delay-Dependent Working Memory Distortions in People With Schizophrenia and Healthy Control Participants. Biol. Psychiatry Cogn. Neurosci. Neuroimaging https://doi.org/10.1016/j.bpsc.2023.07.004 (2023) doi:10.1016/j.bpsc.2023.07.004.

47. Bach, M. The Freiburg Visual Acuity Test-automatic measurement of visual acuity. Optom. Vis. Sci. 73, 49–53 (1996).

48. Houborg, C., Kristjánsson, Á., Tanrikulu, Ö.D. & Pascucci, D. The role of secondary features in serial dependence. J. Vis. 23, 21 (2023).

49. Raftery, A. E. Bayesian model selection in social research. Sociol. Methodol. 111–163 (1995).

50. Brainard, D. H. The psychophysics toolbox. Spat. Vis. 10, 433–436 (1997).

51. Delorme, A. & Makeig, S. EEGLAB: an open source toolbox for analysis of single-trial EEG dynamics including independent component analysis. J. Neurosci. Methods 134, 9–21 (2004).

52. Bigdely-Shamlo, N., Mullen, T., Kothe, C.Su, K.-M. & Robbins, K. A. The PREP pipeline: standardized preprocessing for large-scale EEG analysis. Front. Neuroinformatics 9, 16 (2015).

53. Lehmann, D. & Skrandies, W. Reference-free identification of components of checkerboard-evoked multichannel potential fields. Electroencephalogr. Clin. Neurophysiol. 48, 609–621 (1980).

54. Maris, E. & Oostenveld, R. Nonparametric statistical testing of EEG-and MEG-data. J. Neurosci. Methods 164, 177–190 (2007).

55. Brouwer, G. J. & Heeger, D. J. Decoding and reconstructing color from responses in human visual cortex. J. Neurosci. 29, 13992–14003 (2009).

56. Sprague, T. C. et al. Inverted Encoding Models Assay Population-Level Stimulus Representations, Not Single-Unit Neural Tuning. eNeuro 5, ENEURO.0098-18.2018 (2018).

